# *C. elegans* pathogenic learning confers multigenerational pathogen avoidance

**DOI:** 10.1101/476416

**Authors:** Rebecca S. Moore, Rachel Kaletsky, Coleen T. Murphy

## Abstract

The ability to pass on learned information to progeny could present an evolutionary advantage for many generations. While apparently evolutionarily conserved^1–12^, transgenerational epigenetic inheritance (TEI) is not well understood at the molecular or behavioral levels. Here we describe our discovery that *C. elegans* can pass on a learned pathogenic avoidance behavior to their progeny for several generations through epigenetic mechanisms. Although worms are initially attracted to the gram-negative bacteria *P. aeruginosa* (PA14)^13^, they can learn to avoid this pathogen^13^. We found that prolonged PA14 exposure results in transmission of avoidance behavior to progeny that have themselves never been exposed to PA14, and this behavior persists through the fourth generation. This form of transgenerational inheritance of bacterial avoidance is specific to pathogenic *P. aeruginosa*, requires physical contact and infection, and is distinct from CREB-dependent long-term associative memory and larval imprinting. The TGF-β ligand *daf-7*, whose expression increases in the ASJ upon initial exposure to PA14^14^, is highly expressed in the ASI neurons of progeny of trained mothers until the fourth generation, correlating with transgenerational avoidance behavior. Mutants of histone modifiers and small RNA mediators display defects in naïve PA14 attraction and aversive learning. By contrast, the germline-expressed PRG-1/Piwi homolog^15^ is specifically required for transgenerational inheritance of avoidance behavior. Our results demonstrate a novel and natural paradigm of TEI that may optimize progeny decisions and subsequent survival in the face of changing environmental conditions.

---

*C. elegans* are exposed to and consume a variety of bacterial food sources in their natural environment^16, 17^. Several of these bacteria are pathogens that reduce *C. elegans* lifespan^16^ and progeny production^18, 19^, impacting fitness. Perhaps for this reason, *C. elegans* has evolved the ability to sense its environment and to choose between bacterial food sources. Naïve *C. elegans* initially prefer pathogenic *Pseudomonas aeruginosa* (PA14) to a laboratory strain of nonpathogenic *E. coli* (OP50)^13^. Upon brief exposure to PA14, however, worms rapidly switch their preference and learn to avoid PA14^13^. We wondered whether *C. elegans* can pass this learned avoidance behavior to their naïve progeny (Figure 1A). To test this hypothesis, we exposed wild-type L4 hermaphrodites to PA14 for 4 h (as in previous training regimens^13^) and bleached the mothers to obtain F1 eggs, but we found that this short exposure does not induce transgenerational effects in the adult progeny (Figure 1A-B). By contrast, when we exposed the mothers to PA14 for 24 h, the adult F1 progeny exhibited avoidance, despite never previously having encountered PA14 (Figure 1C).

**Figure 1:**
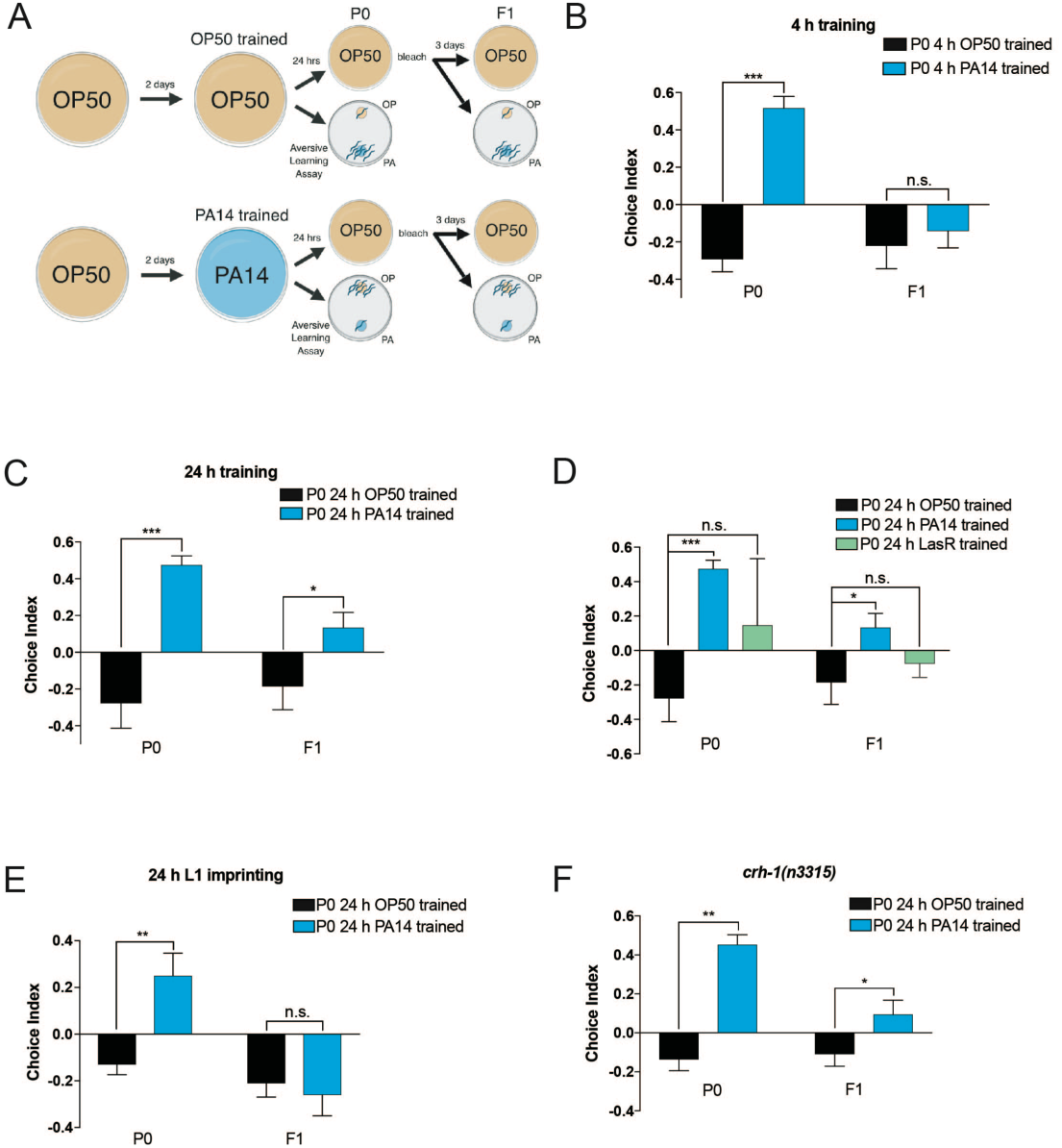
Pathogenic avoidance is transgenerationally inherited in *C. elegans*. (A) Adult pathogen training protocol: wild-type worms were bleached onto OP50 (OP) *E.* coli plates; two days later, L4 stage worms were transferred to OP50 or *Pseudomonas aeruginosa* PA14 (PA) training plates that have been incubated for two days at 25°C. Worms remained on training plates for 24 h at 20°C. Immediately following training, worms were subjected to an aversive learning assay (modified from Zhang et al., 2005)^13^ or bleached onto OP50-seeded plates. F1 animals were tested as Day 1 adults. (B) 4 h of training on PA14 is sufficient to elicit maternal avoidance (P0) of PA14, but is not sufficient for progeny avoidance (F1). Choice Index = (# of worms on OP50 - # of worms on PA14)/(Total # of worms) (C) 24 h of training on PA14 induces both pathogen avoidance and transgenerational inheritance of pathogen avoidance (F1). (D) 24 h of training on avirulent Pseudomonas (LasR) does not induce maternal pathogenic learning or progeny avoidance of PA14. (E) L1-imprinted animals (24 h) exhibit adult (Day 1) aversion to PA14, but progeny of imprinted mothers do not. (F) CREB (*crh-1*) is not required for maternal pathogenic learning or progeny avoidance of PA14. One Way ANOVA, Tukey’s multiple comparison test, mean ± SEM. n ≥ 7-10 per generation (n = one choice assay plate with 50-200 worms per plate). *p ≤ 0.05, **p ≤ 0.01, ***p ≤ 0.001, ns = not significant. At least 3 biological replicates were performed for all aversive learning assays.

A previous study reported that exposure to the odor of PA14 or a chemical, 2-aminoacetophenone (2AA), can cause PA14 avoidance^20^. We wondered whether training with odor alone would be sufficient to induce transgenerational memory. However, in our hands, neither mothers nor progeny of mothers exposed to PA14 odor or 2AA avoided PA14 (Supplement 1A-B), suggesting that physical contact is required for both pathogenic learning and for the transgenerational inheritance of this behavior. Next we asked whether virulence is required for these effects. Prolonged PA14 exposure kills *C. elegans* within 60 hrs^18^, cutting short their normal 2-3 week lifespan (Supplement 1C). The PA14 quorum-sensing mutant LasR is markedly less virulent to *C. elegans*, such that LasR-exposed worms remain alive after ~2.5 days exposure when all of the wild-type PA14-exposed animals have died^18^. We found that wild-type worms exposed to LasR do not learn to avoid PA14 (Figure 1D), and progeny of LasR-trained mothers also fail to avoid PA14 (Figure 1D). Together, these results suggest that pathogenic learning and transgenerational inheritance of pathogen avoidance require both physical contact and infection with PA14.

Early developmental stage larvae (L1) are capable of learning to avoid PA14^21^. This process, called imprinting, results in the maintenance of PA14 avoidance in early adulthood, but is not transmitted to progeny of imprinted mothers^21^. Because imprinting training is typically 12 hours^21^, we asked whether longer L1 training would cause both adult aversion to PA14 and TEI of pathogen avoidance behavior. Although 24 h of larval training was sufficient to elicit parental avoidance of PA14 (Figure 1E), progeny of parents trained as L1s did not avoid PA14 (Figure 1E). Furthermore, the transcription factor CREB (*crh-1*), which is required for L1 imprinting of PA14^21^ and for many forms of longterm memory^22^, is not necessary for transgenerational pathogenic avoidance, further distinguishing these processes (Figure 1F). Therefore, the mechanism of TEI is distinct from other forms of CREB-dependent, aversive long-term memory^22^, including larval imprinting of PA14 avoidance^21^.

We next wanted to determine the number of generations this avoidance lasts. During training, progeny themselves may have been exposed to the bacterial pathogen while still in the mother, in which case the F1 avoidance effect could not be considered truly transgenerational, but such an effect should be restricted to one generation. However, we found that training of mothers (P0) induces pathogenic avoidance in the four subsequent generations, F1-F4 (Figure 2A-D; F). Fifth generation (F5) descendants no longer avoid PA14 but instead are attracted to PA14, resuming naïve behavior (Figure 2E; F).

**Figure 2:**
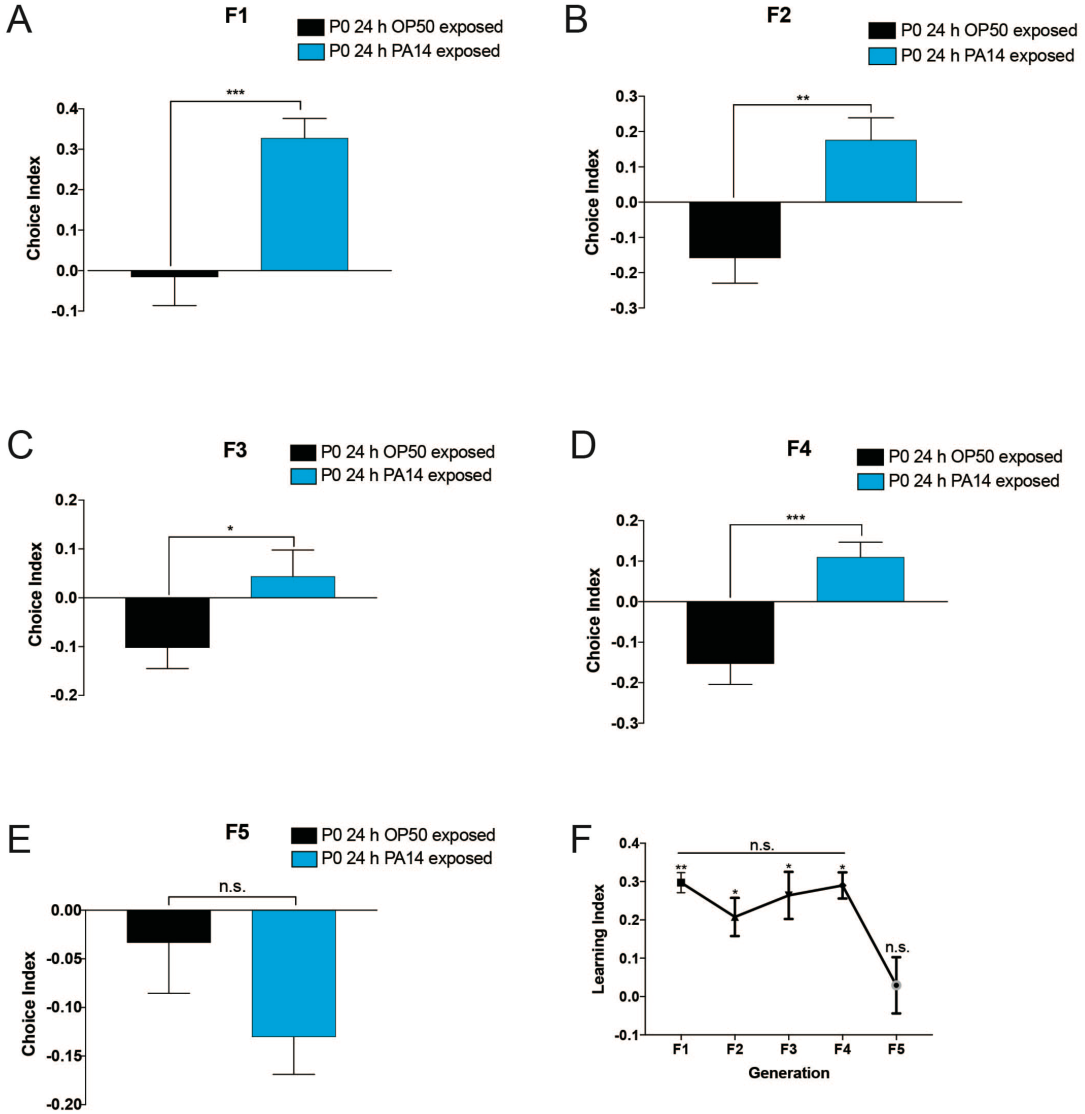
Transgenerational inheritance of PA14 is multigenerational. (A-E) Untrained (naïve) progeny of PA14- trained mothers avoid PA14 from generation F1 through F4. The fifth generation returns to normal PA14 attraction (E). Students t-test, mean ± SEM. n = 10 per generation (n = one choice assay plate with 50-200 worms per plate). (F) Learning index (naïve choice index – trained choice index) of generation F1 through F5. One Way ANOVA, Tukey’s multiple comparison test, mean ± SEM. n ≥ 7-10 per generation. *p ≤ 0.05, **p ≤ 0.01, ***p ≤ 0.001, ns = not significant. At least 3 biological replicates were performed per generation for all aversive learning assays.

Sensing and subsequent avoidance of *Pseudomonas* requires the activity of the nervous system^13, 14, 19, 23, 24^, and we wondered whether transgenerational avoidance utilizes known components of PA14 sensing. Meisel, et al. (2014)^14^ previously showed that the TGF-β ligand DAF-7 is expressed basally in ASI sensory neurons, and upon PA14 exposure, *daf-7* expression increases in the ASI and is induced in the ASJ neurons. (DAF-7 signaling in the ASJ controls expression of TGF-β signaling in downstream RIM/RIC interneurons, which in turn control reversals through downstream motor neurons^23^) We confirmed these observations in mothers exposed to PA14 for 24 h (Figure 3A-D). We then asked whether transgenerational avoidance behavior training induces higher expression of *daf-7::gfp* in the ASI and ASJ in progeny of trained mothers. Surprisingly, F1 progeny of PA14-trained mothers maintained a high level of *daf-7::gfp* expression in the ASI (Figure 3E-G), while *daf-7::gfp* expression in the ASJ returned to basal levels in these naïve progeny (Figure 3F). However, upon brief exposure to PA14, *daf-7::gfp* expression in the ASJ was induced to higher levels in the progeny of PA14 trained-mothers compared to OP50 trained-mothers within 4 hours (Figure 3H). Because the choice assay measures an animal’s food preference within a short time frame (<1 h), our *daf-7::gfp* expression data suggest that the basal elevation of *daf-7* in the ASI of progeny, rather than its delayed induction in the ASJ, mediates transgenerational avoidance behavior. We then asked whether the ASI is necessary for TEI of pathogen avoidance. Mutants in which the ASI is genetically ablated avoid PA14 after training, similar to wild-type animals (Figure 3I). However, progeny of pathogen-trained mutants do not avoid PA14 (Figure 3J), indicating that the ASI is required for TEI of pathogen avoidance. Thus, while the ASI and ASJ are redundant for pathogenic learning^14^, the ASI is required for transgenerational inheritance of the pathogenic avoidance behavior. As we found for pathogenic learning, training with avirulent LasR *Pseudomonas* did not increase *daf-7::gfp* expression in the ASI, nor did it induce expression of *daf-7::gfp* in the ASJ of mothers or their progeny (Figure 3C, G), suggesting that virulence is required for transgenerational *daf-7::gfp* expression changes.

**Figure 3:**
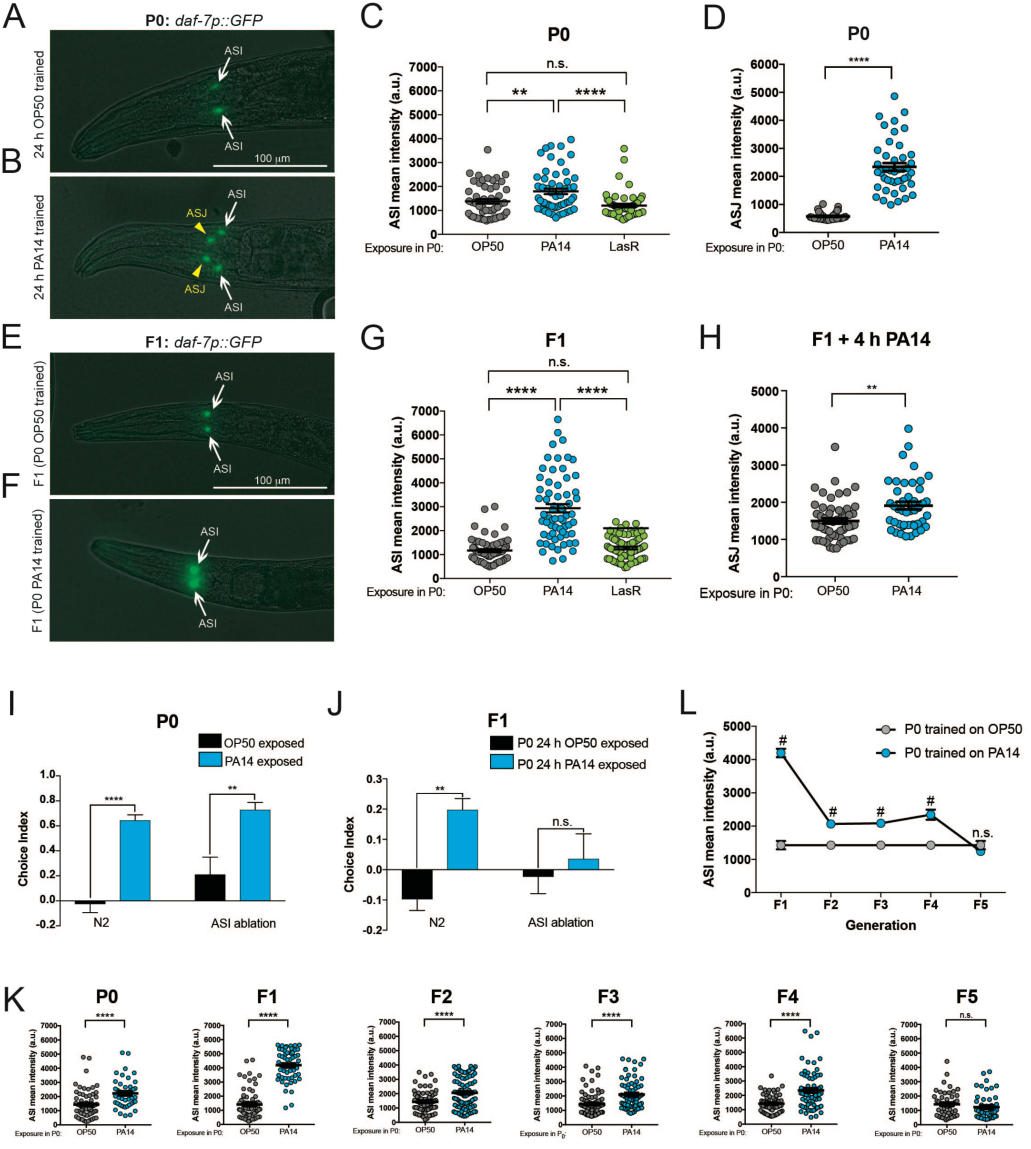
*daf-7::gfp* expression remains elevated F1-F4 progeny of pathogen-exposed mothers. (A) *daf-7::gfp* is expressed in the ASI neuron of naïve animals (white arrow). (B, D) PA14 training induces *daf-7::gfp* expression in the ASJ neuron (yellow arrow head). Students t-test, mean ± SEM. n ≥ 36-43 per training condition (n = individual neuron, minimum of 18 worms). At least 3 biological replicates were performed for all assays. (C) PA14 training increases *daf-7::gfp* expression in the ASI, compared to training with OP50 or LasR (avirulent PA14 mutant). One Way ANOVA, Tukey’s multiple comparisons test, mean ± SEM. n ≥ 54-56 per training condition (n = individual neuron, minimum of 27 worms). At least 2 biological replicates were performed for all assays. (E-G) Naïve progeny of PA14-trained mothers increase expression of *daf-7::gfp* in the ASI compared to progeny of OP50- or LasR-trained progeny. One Way ANOVA, Tukey’s multiple comparisons test, mean ± SEM. n ≥ 61-75 per training condition (n = individual neuron, minimum of 32 worms). At least 2 biological replicates were performed for all assays. (H) Progeny (F1) of PA14-trained mothers have higher *daf-7::gfp* expression in the ASJ after 4 h of PA14 training. Students t-test, mean ± SEM. n ≥ 44-55 per training condition (n = individual neuron, minimum of 18 worms). (I) The ASI is not required for maternal (P0) pathogenic learning. One Way ANOVA, Tukey’s multiple comparison test, mean ± SEM. n ≥ 7-10 per generation (n = one choice assay plate with 50-200 worms per plate). *p ≤ 0.05, **p ≤ 0.01, ***p ≤ 0.001, ns = not significant. (J) The ASI is required for progeny (F1) avoidance of PA14. One Way ANOVA, Tukey’s multiple comparison test, mean ± SEM. n ≥ 7-10 per generation (n = one choice assay plate with 50-200 worms per plate). *p ≤ 0.05, **p ≤ 0.01, ***p ≤ 0.001, ns = not significant. (K) PA14 training increases *daf-7::gfp* expression in the ASI of progeny of PA14-trained mothers for 4 generations (F4) before returning to low levels in the 5^th^ generation (F5), following a similar trajectory as avoidance behavior (L) # = ****p < 0.0001. Students t-test, mean ± SEM. n ≥ 44-98 per training condition per generation (n = individual neuron, minimum of 22 worms). 2 biological replicates were performed. *p ≤ 0.05, **p ≤ 0.01, ***p ≤ 0.001, ****p < 0.0001, ns = not significant.

Next, we asked how many generations the increased expression of ASI *daf-7::gfp* persists after pathogenic training. ASI *daf-7::gfp* remained elevated in progeny of PA14 trained-mothers for four generations (Figure 3K, L), then returned to basal levels in the F5 (Figure 3K, L), similar to the trajectory of transgenerational avoidance behavior in the F1-F4 generations (Figure 2F). Together, these results suggest that progeny who inherit PA14 avoidance behavior maintain high ASI *daf-7::gfp* expression levels, rendering the animals “primed” for increased expression of *daf-7::gfp* in the ASJ and subsequent avoidance upon PA14 encounter, and that ASI *daf-7* levels set the avoidance response ability of that generation.

To understand the mechanisms underlying transgenerational pathogenic avoidance, we examined the role of candidate TEI regulators, such as histone^1–5^, siRNA^3, 6–8^, and piRNA^9–12^ modifiers. Transient mutation of the COMPASS histone modification complex components, which are expressed and function in the germline^25^, induce a transgenerational longevity effect that, like TEI of pathogen avoidance, lasts for 4 generations^4^. However, we found that COMPASS complex mutants *set-2*, an H3K4me3 methyltransferase, and *rbr-2*, an H3K4me3 demethylase, were already defective for naïve PA14 attraction, so their effect on aversive learning or subsequent transgenerational effects is unclear (Figure 4A, Supplement 2A, 2C). Similarly, mutants of *set-32*, a putative H3K9me3 methyltransferase that links histone modifications and siRNAs^1^ in transgenerational signaling, were defective in pathogenic learning (Figure 4B), as were progeny of PA14 trained mothers (Supplement 2B, 2D).

**Figure 4:**
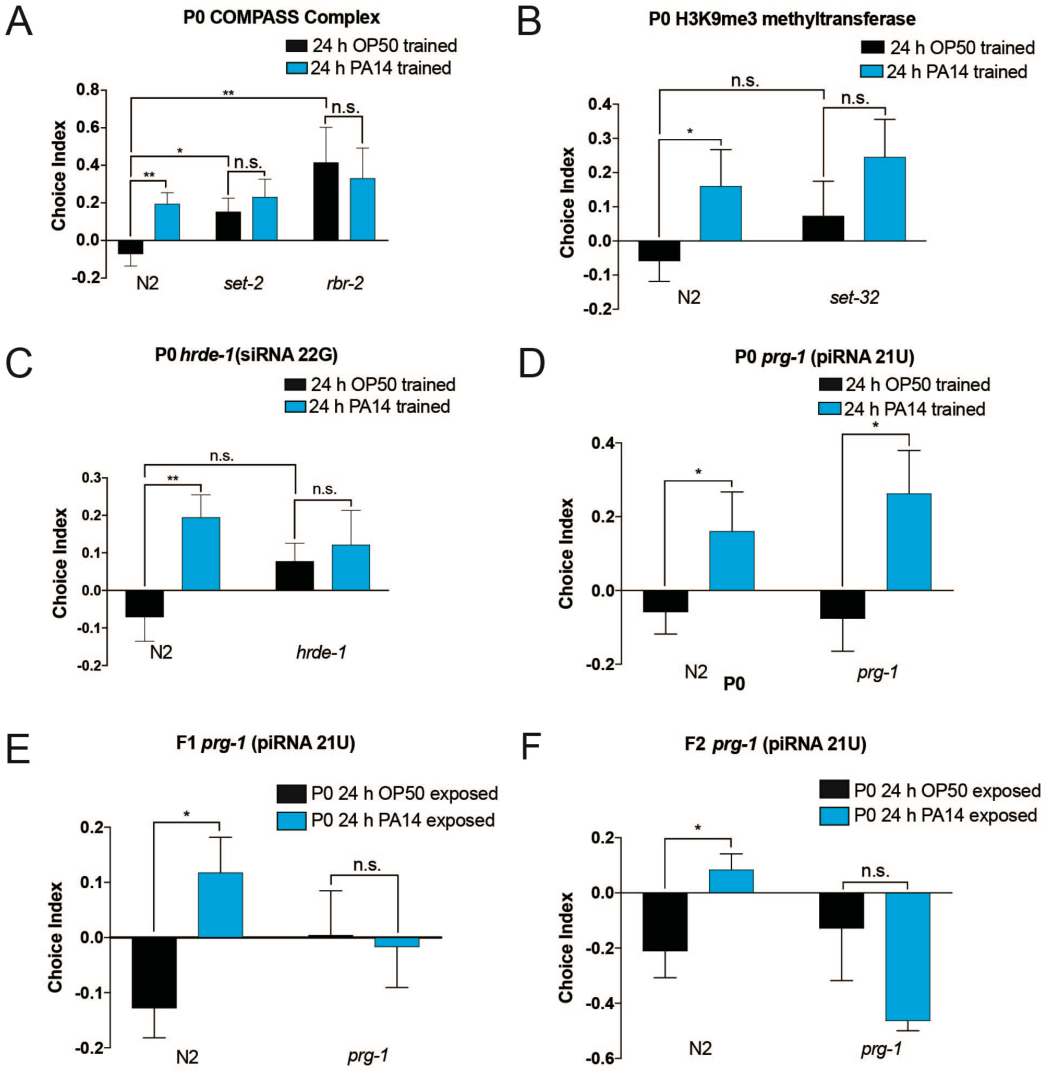
PRG-1/Piwi is required for transgenerational inheritance of pathogenic aversive learning. (A) COMPASS complex mutants (*set-2, rbr-2*) are defective in naïve PA14 preference. (B) *set-32* (H3K9 methyltransferase) mutants have normal naïve PA14 preference, but are defective for pathogenic learning. (C) *hrde-1* mutants have normal naïve preference, but are defective for aversive learning after training on PA14. (D) Like wild-type, *prg-1* mutants have normal naïve preference and can learn to avoid PA14 after training. (E) Unlike wild-type, naive progeny (F1) of PA14-trained progeny do not avoid PA14. (F) Naïve *prg-1* F2 (grandprogeny) of PA14-trained mothers do not avoid PA14. One Way ANOVA, Tukey’s multiple comparison test, mean ± SEM. n ≥ 6-10 per generation. *p ≤ 0.05, **p ≤ 0.01, ns = not significant. At least 2 biological replicates were performed per generation for all aversive learning assays.

Small RNA regulators, such as the nuclear argonaute HRDE- 1/Ago and the PIWI argonaute PRG-1, are involved in initiation^26^ and transmission^7^ of small RNAs, respectively, but whether they play a role in behavior has not yet been demonstrated. Like *set-32* and the COMPASS mutants, *hrde-1* mutants were defective for pathogenic aversive learning (Figure 3C, Supplement 2E). By contrast, mutants of *prg-1/Piwi* display normal naïve choice preference and pathogenic learning after PA14 training (Figure 4D), but the progeny of trained *prg-1/Piwi* mothers were defective in their avoidance of PA14 (Figure 4E, 4F), suggesting that PRG-1/Piwi is required for transgenerational avoidance behavior.

Defects in naïve attraction and pathogenic learning in the P0 complicate the study of transgenerational effects. To specifically test whether the COMPASS complex, *set-32*, or *hrde-1* are involved in transgenerational memory, we first carried out P0 training on wild-type animals (Supplement 3A), and then used RNAi to knock down COMPASS complex components/*set-32/hrde-1* in the F1 generation. Although the mothers of these progeny had learned to avoid PA14 (Supplement 3A), RNAi treatment of progeny of PA14-trained mothers abolished the learning effect, resulting in a return to naïve preference (Supplement 3B-E). Thus, *set-2*, *rbr-2*, *set-32*, and *hrde-1* are required for the processes of sensing pathogenic PA14 and learning pathogenic avoidance, and *set-2* and *set-32* are further required in the F1 for expression of the avoidance behavior. In contrast, PRG-1/Piwi function is not required for initial avoidance learning, but is specifically necessary for transmission of the learned behavior from mothers to progeny.

Transgenerational pathogen avoidance and multi-generational longevity after transient loss of COMPASS complex components share similar kinetics, functioning in the F1 through F4, but not the F5 generation (Figure 2F). Because we were able to observe this avoidance behavior in wild-type animals rather than in an artificial (transient mutant) situation, this transgenerational pathogenic avoidance paradigm may represent the natural context for such behaviors. Indeed, naïve progeny of PA14-trained mothers survive longer on a small PA14 lawn (where there is an opportunity to avoid the pathogen) compared to OP50-trained controls (Figure 5A). The increased survival is dependent on the ability to avoid PA14 on the assay plates, since obligate PA14 exposure confers no survival difference (Figure 5A).

**Figure 5:**
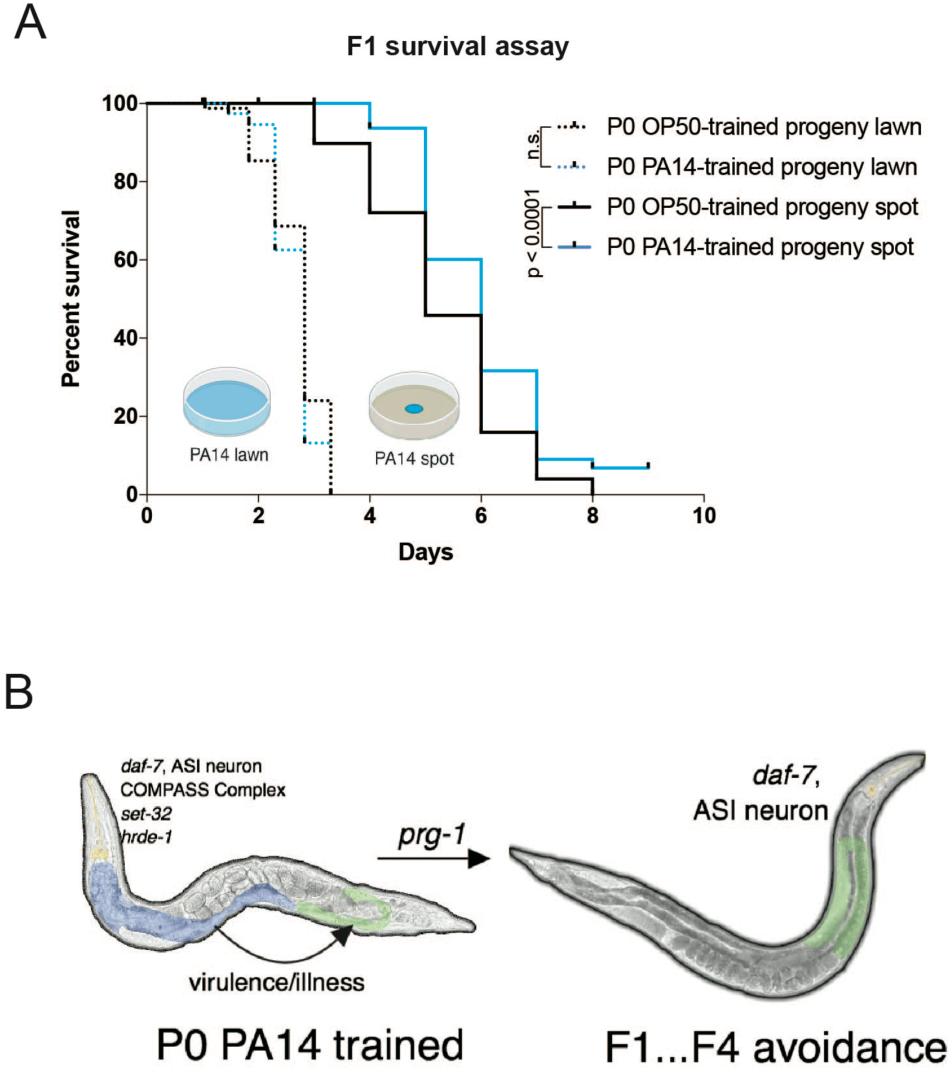
Transgenerational inheritance of avoidance behavior confers a survival benefit to *C. elegans* progeny. (A) Progeny of PA14-trained mothers and progeny of OP50-trained mothers have a similar survival span on a full lawn of PA14 (p = 0.2931). However, progeny of PA14-trained mothers have a survival advantage on a spot of PA14 compared to progeny of OP50-trained mothers (p < 0.0001). Log-rank (Mantel-Cox) test. n ≤ 80-120 worms per condition. (B) MODEL: Upon interaction with PA14, *C. elegans* sense PA14 and learn to avoid PA14 upon subsequent exposure, which requires the *daf-7* in the ASI neuron, COMPASS complex, SET-32, and HRDE-1. Once pathogen avoidance has been learned, a signal is sent through PRG-1 to induce transgenerational inheritance of pathogenic avoidance from F1 through F4 generations. Yellow = neurons, Blue = intestine, Green = germline.

Several strategies are employed by animals exposed to pathogens to maintain fitness in the face of imminent death. *C. elegans* directly exposed to PA14 immediately engage innate immune responses to combat the pathogen^27^, and subsequently induce behavioral avoidance to avoid the pathogen^13^

Here, we describe a new behavioral component of the pathogen response, whereby naïve progeny avoid PA14 as a consequence of the memory of pathogen exposure in previous generations (Figure 5B). *Pseudomonas* species (including non-pathogenic *Pseudomonas*) comprise up to 30% of the bacteria in *C. elegans’* natural environments^16, 17^ and thus may provide a substantial nutritional source. Therefore, avoidance of all *Pseudomonas* after pathogen exposure might be a poor long-term strategy. Instead, temporary avoidance of pathogenic *Pseudomonas* might drive the worms’ progeny to escape pathogenic food sources, while eventually allowing the return to a potentially nutritious *Pseudomonas* species for food. This likely serves to increase the fitness of subsequent generations in the response to prolonged pathogen exposure that may be encountered in the wild. Together, our results demonstrate that transgenerational avoidance of pathogenic bacteria provides a biological context for TEI in *C. elegans*, where animals must distinguish between both beneficial and detrimental food sources to ensure survival of self and progeny in a dynamic environment.

## Methods

### *C. elegans* and bacterial strains and cultivation

Strains were provided by the CGC. SX922: *prg-1(n4357)*, RB1025: *set-2 (ok952)*, ZR1: *rbr-2(tm1231)*, VC967: *set-32(ok1457)*, FK181: *ksIs2 [Pdaf-7::GFP + rol-6(su1006)]*, MT9973: *crh-1(n3315)*, PY7505: oyIs84 [*gpa-4p*::TU#813+*gcy-27p*::TU#814 + *gcy-27p*::GFP + *unc-122p*::DsRed]. Strains were provided by the National Bioresource *hrde1(tm1200)*. OP50 and OP50-1 were provided by the CGC. PA14 and LasR were gifts from Z. Gitai. Worm strains were maintained at 15°C on HGM plates on *E. coli* OP50 using standard methods.

### Pathogen training

Eggs from young adult hermaphrodites were obtained by bleaching and placed on to High Growth (HG) plates and left at 20°C for 2 days. Training plates were prepared by inoculating overnight cultures of OP50 and pathogen in LB at 37°C. Overnight cultures were diluted in LB to an Optical Density (OD_600_) = 1 and used to seed Nematode Growth Media (NGM) plates. Plates were incubated at 25°C in separate incubators for 2 days. On day of training (i.e., 2 days post bleaching) plates were left to cool on a bench top for < 1 hr. 10 μL of pooled L4 worms were plated onto OP50 seeded training plates, while 40 μL of worms were plated onto pathogen seeded training plates. Worms were incubated on training plates at 20°C in separate containers for 24 hrs. After 24 hrs, worms were washed off plates in M9 3x. Some worms were used for an aversive learning assay, while the majority of worms were bleached onto HG plates at 20°C for 3 days. For experiments involving RNAi; animals were trained using pL4440 empty vector as control RNAi in HT115 bacteria.

### Aversive learning assay

Overnight bacterial cultures were diluted in LB to an Optical Density (OD_600_) = 1, and 25 μL of each bacterial suspension was seeded on a 60 mm NGM plate and incubated at 25°C for 2 days. After 2 days assay plates were left at room temperature for 1 hr before use. Immediately before use, 1 μL of 1M sodium azide was spotted onto each bacterial lawn to be used as a paralyzing agent during choice assay. To start the assay (modified from Zhang et al., 2005), worms were washed off training plates in M9, and washed 2 additionally times in M9. 5 μL of worms were spotted at the bottom of the assay plate, using a wide orifice tip, midway between the bacterial lawns. Assays were incubated at room temperature for 1 hr before counting the number of worms on each lawn.

In experiments in which F1 and subsequent generations are used: All animals tested are washed off HG plates with M9 at Day 1. Some of the pooled animals are subjected to an aversive learning assay, while the majority of worms are bleached onto HG plates left at 20°C for 3 days and used to test F2s.

### Odor + 2-aminoacetophenone (2AA) training

3 mL of fresh overnight bacterial cultures (OD_600_ OP50 = 1.4- 1.7, OD_600_ PA14 = 3.3-3.5), water, or 2AA (catalog no. A38207, Sigma-Alderich) (1 mM, diluted in water) were placed into 2 lids of a 35 mm petri dish, respectively, which was then placed in the lid of an inverted 10 cm NGM petri dish that had been prepared as described for pathogen training. Odor training assays were left in the dark at room temperature for 24 hrs. All training conditions were maintained in separate containers.

### L1 imprinting

Eggs from young hermaphrodites were obtained by bleaching and placed directly onto OP50 or pathogen prepared training plates (5 μL of eggs were placed onto OP50 training plates, and 20 μL of eggs were placed onto PA14 training plates). Plates were incubated at 20°C for 24 hrs. After 24 hrs, worms were washed off training plates using M9 + 50 mg/mL streptomycin. Worms were washed 2 times and plated onto HGM+300 mg/mL streptomycin plates seeded with OP50-1 (streptomycin resistant OP50). Worms were left to mature to Day 1 adults and used in an aversive learning assay. Subsequent generations were prepared by bleaching pooled animals onto HG plates.

### PA14 survival assay

1 mL of fresh overnight bacterial cultures was diluted in 4 mL of LB. For full lawn assays 750 μL of diluted PA14 was spread to completely cover a 10 cm NGM plate. For PA14 spot assays, 100 μL of diluted PA14 was placed in the center of a 10 cm NGM plate resulting in a 2 cm spot in the center of the pate. Plates were incubated for 2 days at 25°C before use. Upon addition of Day 1 worms to plates, assays were performed at 20°C. PA14-lawn assays were counted every 6-8 h. PA14-spot assays were counted every 24 h. Every 48 h, worms in both assays were moved onto new plates. For spot assays, animals were transferred to similar locations on new plates.

### RNAi Interference

RNAi experiments were conducted using the standard feeding RNAi method. Bacterial clones expressing the control (empty vector, pL4440) construct and the dsRNA targeting *C. elegans* genes were obtained from the Ahringer RNAi library. All RNAi clones were sequenced prior to use. RNAi-induced knockdown was conducted by bleaching progeny of HT115 pL4440 or PA14 trained animals onto RNAi seeded plates.

### Imaging and fluorescence quantitation

Z-stack multi-channel (DIC, GFP) of day 1 adult GFP transgenic worms were imaged every 1 μm at 60X magnification; Maximum Intensity Projections and 3D reconstructions of head neurons were built with Nikon *NIS-Elements*. To quantify *daf-7p*::GFP levels, worms were prepared and treated as described for pathogen training. Worms were mounted on agar pads and immobilized using 1 mM levamisole. GFP was imaged at 60X magnification and quantified using *NIS Elements* software. Average pixel intensity was measured in each worm by drawing a bezier outline of the neuron cell body for 2 ASI head neurons and/or 2 ASJ head neurons.

## Acknowledgements

We thank the *C. elegans* Genetics Center for strains; WormBase (WS250); Z. Gitai for bacteria strains, and the Murphy lab for discussion. C.T.M. is the Director of the Glenn Center for Aging Research at Princeton. R.S.M. was funded by T32GM007388 (NIGMS), and further support was provided by a DP1 Pioneer Award to C.T.M.

## Author contributions

R.S.M., R.K., and C.T.M. designed experiments. R.S.M. and R.K. performed experiments and analyzed data. R.S.M., and R.K. performed aversive learning and chemotaxis experiments. R.K. performed imaging and quantification experiments. R.S.M., R.K., and C.T.M wrote the manuscript.

---

**Supplement 1:**
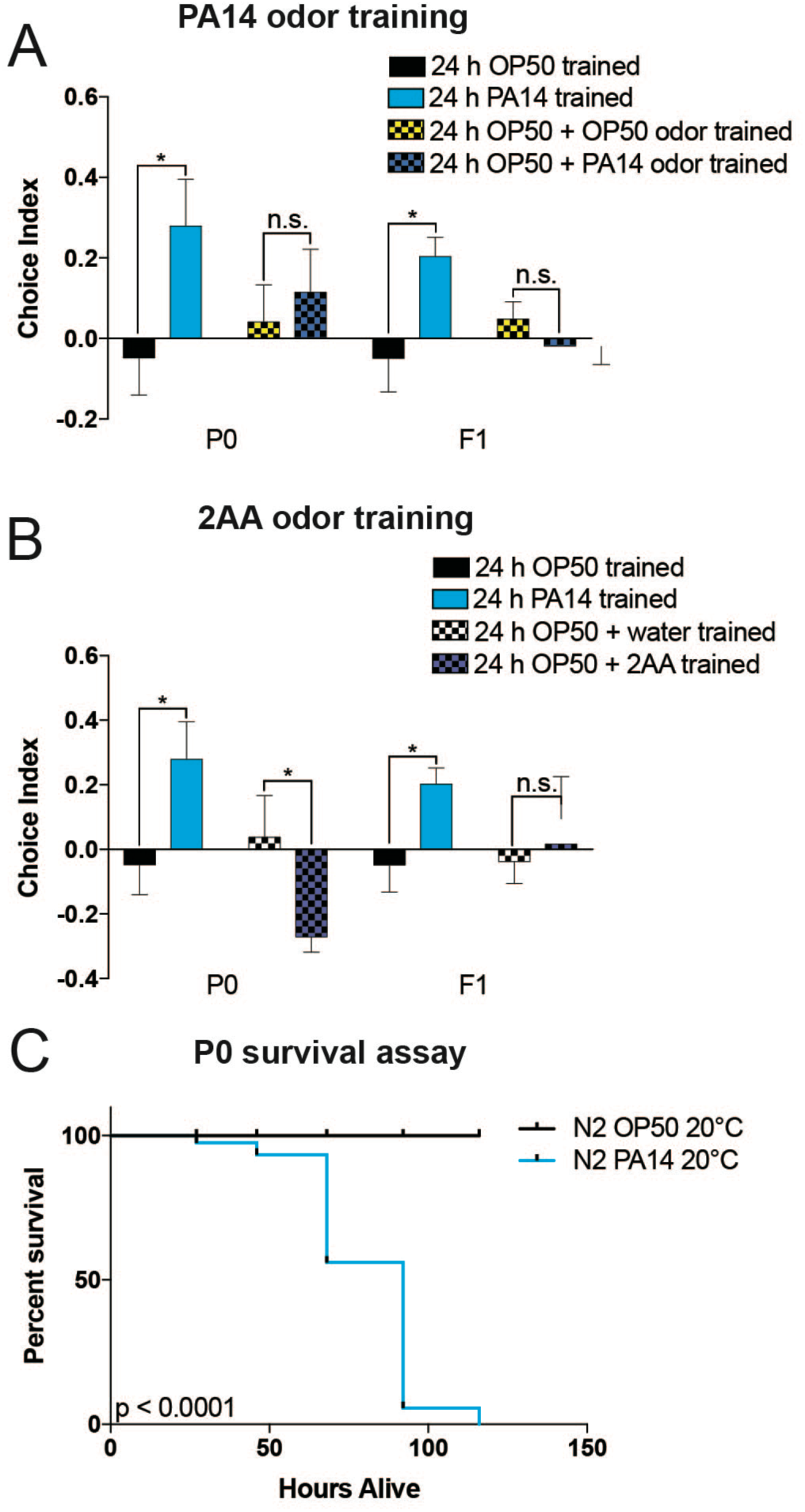
Transgenerational inheritance of pathogen avoidance requires direct contact with PA14. (A) PA14 odor is not sufficient to induce aversive pathogenic learning. (B) 2-aminoacetophenone (2AA) is not sufficient to induce aversive pathogenic learning compared to water alone. One Way ANOVA, Tukey’s multiple comparison test, mean ± SEM. n ≥ 6-10 per generation (n = one choice assay plate with 50-200 worms per plate). *p ≤ 0.05, **p ≤ 0.01, ns = not significant. At least 2 biological replicates were performed for all aversive learning assays. (C) PA14 is lethal at 20°C. p < 0.0001, Log-rank (Mantel-Cox) test. n ≥ 78-81 worms per condition.

**Supplement 2:**
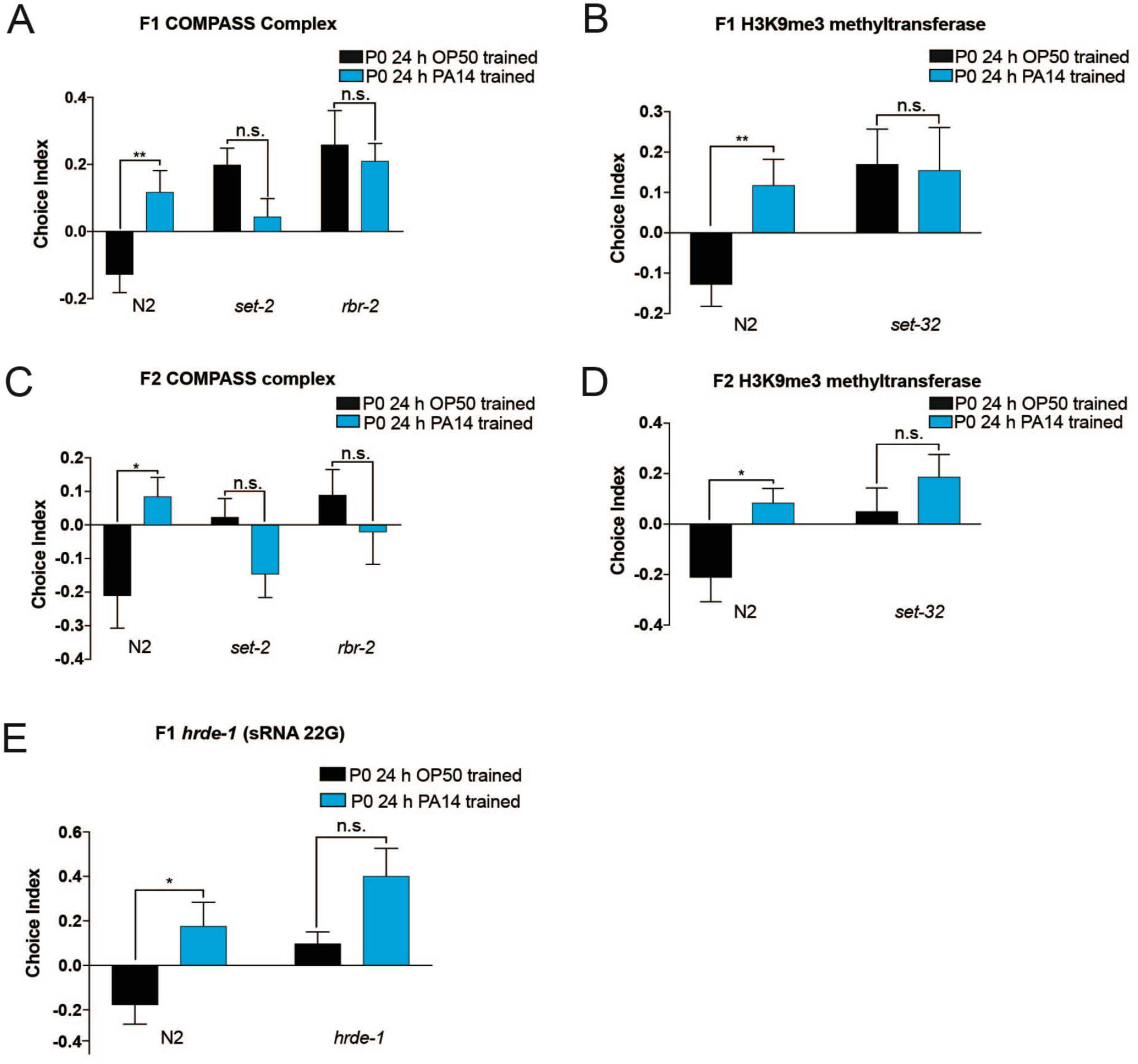
Mutants that do not learn avoidance behavior exhibit similar behavior in F1 and F2 following P0 training. (A-B) F1 progeny of PA14-trained *set-2, rbr-2*, and *set-32* are defective in pathogenic-aversive learning. (C-D) F2 progeny of PA14-trained *set-2*, *rbr-2*, and *set-32* are defective in pathogenic-aversive learning. (E) Progeny of *hrde-*1 PA14-trained mutants are defective in pathogenic-aversive learning. One Way ANOVA, Tukey’s multiple comparison test, mean ± SEM. n ≥ 6-10 per generation (n = one choice assay plate with 50-200 worms per plate). *p ≤ 0.05, **p ≤ 0.01, ns = not significant. At least 2 biological replicates were performed for all aversive learning assays.

**Supplement 3:**
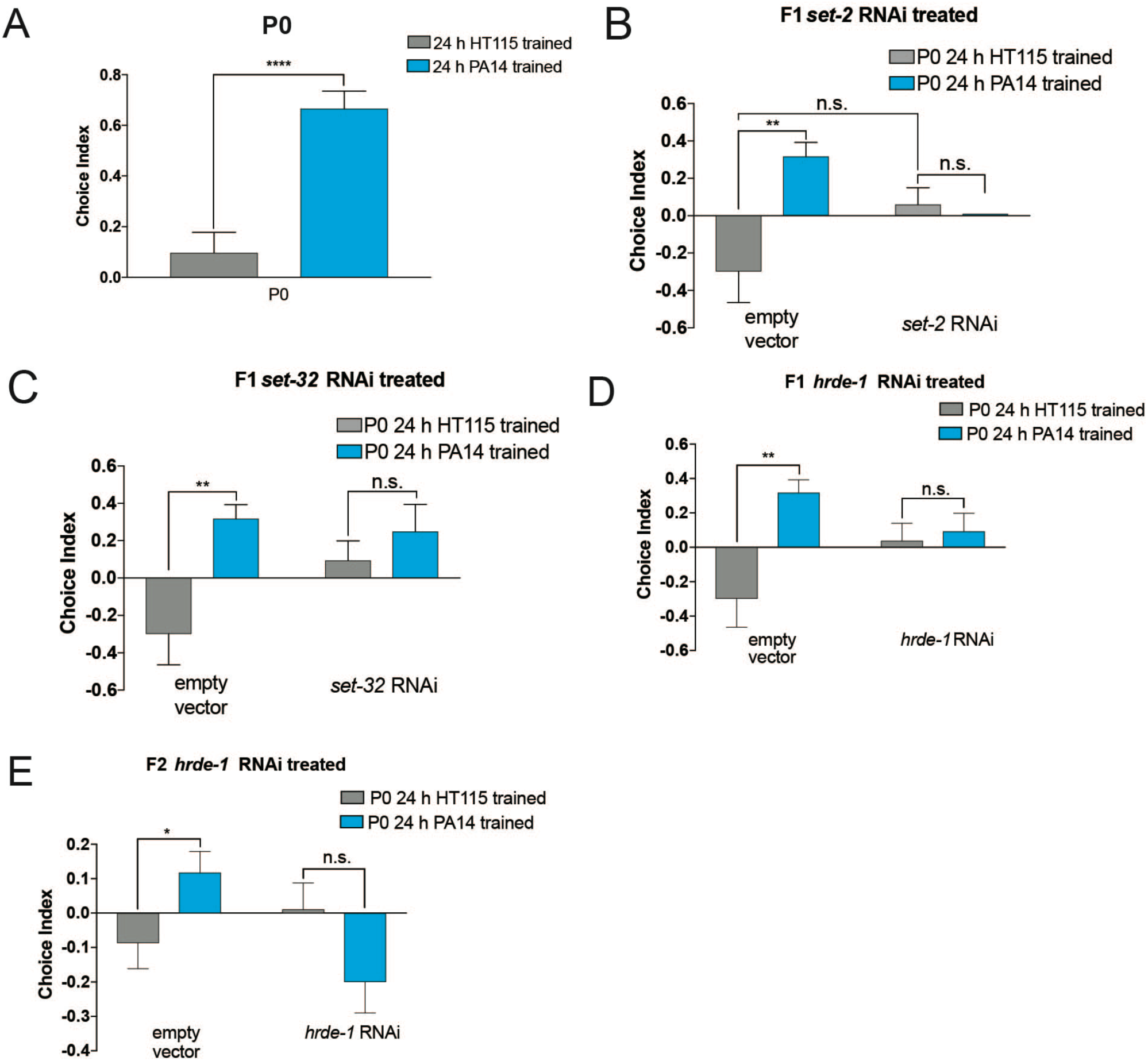
RNAi-treated progeny of PA14-trained mothers still exhibit aversive learning defects in the F1 and F2 generations. (A) 24 h of PA14 training results in aversive learning compared to 24 h of training with *E. coli* HT115. (B) F1 progeny of PA14-trained mothers treated with RNAi of *set-2* were defective in pathogen learning. (C) F1 Progeny of PA14- trained mothers treated with RNAi of *set-32* were defective in pathogen learning. (D-E) F1 and F2 progeny treated with *hrde-1* RNAi of PA14-trained mothers are defective in pathogenic learning. One Way ANOVA, Tukey’s multiple comparison test, mean ± SEM. n = 10 per generation (n = one choice assay plate with 50-200 worms per plate). *p ≤ 0.05, **p ≤ 0.01, ***p ≤ 0.001, ****p < 0.0001, ns = not significant. At least 2 biological replicates were performed for all aversive learning assays.

## References

1. Woodhouse, R. M. et al. The chromatin modifiers SET-25 and SET-32 are required for initiation but not long-term maintenance of transgenerational epigenetic inheritance. (2018). doi:10.1101/255646

2. Klosin, A., Casas, E., Hidalgo-Carcedo, C., Vavouri, T. & Lehner, B. Transgenerational transmission of environmental information in C. elegans. Science 356, 320–323 (2017).

3. Spracklin, G. et al. The RNAi Inheritance Machinery of Caenorhabditis elegans. Genetics 206, 1403–1416 (2017).

4. Greer, E. L. et al. Transgenerational epigenetic inheritance of longevity in Caenorhabditis elegans. Nature 479, 365–371 (2011).

5. Vassoler, F. M., White, S. L., Schmidt, H. D., Sadri-Vakili, G. & Pierce, R. C. Epigenetic inheritance of a cocaine-resistance phenotype. Nat. Neurosci. 16, 42–47 (2013).

6. Rechavi, O. et al. Starvation-Induced Transgenerational Inheritance of Small RNAs in C. elegans. Cell 158, 277–287 (2014).

7. Buckley, B. A. et al. A nuclear Argonaute promotes multigenerational epigenetic inheritance and germline immortality. Nature 489, 447–451 (2012).

8. Rechavi, O., Minevich, G. & Hobert, O. Transgenerational Inheritance of an Acquired Small RNA-Based Antiviral Response in C. elegans. Cell 147, 1248–1256 (2011).

9. Heestand, B., Simon, M., Frenk, S., Titov, D. & Ahmed, S. Transgenerational Sterility of Piwi Mutants Represents a Dynamic Form of Adult Reproductive Diapause. Cell Rep. 23, 156–171 (2018).

10. Ashe, A. et al. piRNAs Can Trigger a Multigenerational Epigenetic Memory in the Germline of C. elegans. Cell 150, 88–99 (2012).

11. Brennecke, J. et al. An Epigenetic Role for Maternally Inherited piRNAs in Transposon Silencing. Science 322, 1387–1392 (2008).

12. Grentzinger, T. et al. piRNA-mediated transgenerational inheritance of an acquired trait. Genome Res. 22, 1877–1888 (2012).

13. Zhang, Y., Lu, H. & Bargmann, C. I. Pathogenic bacteria induce aversive olfactory learning in Caenorhabditis elegans. Nature 438, 179–184 (2005).

14. Meisel, J. D., Panda, O., Mahanti, P., Schroeder, F. C. & Kim, D. H. Chemosensation of Bacterial Secondary Metabolites Modulates Neuroendocrine Signaling and Behavior of C. elegans. Cell 159, 267–280 (2014).

15. Batista, P. J. et al. PRG-1 and 21U-RNAs Interact to Form the piRNA Complex Required for Fertility in C. elegans. Mol. Cell 31, 67–78 (2008).

16. Samuel, B. S., Rowedder, H., Braendle, C., Félix, M.-A. & Ruvkun, G. Caenorhabditis elegans responses to bacteria from its natural habitats. Proc. Natl. Acad. Sci. 113, E3941–E3949 (2016).

17. Dirksen, P. et al. The native microbiome of the nematode Caenorhabditis elegans: gateway to a new host-microbiome model. BMC Biol. 14, (2016).

18. Tan, M. W., Mahajan-Miklos, S. & Ausubel, F. M. Killing of Caenorhabditis elegans by Pseudomonas aeruginosa used to model mammalian bacterial pathogenesis. Proc. Natl. Acad. Sci. U. S. A. 96, 715–720 (1999).

19. Lee, K. & Mylonakis, E. An Intestine-Derived Neuropeptide Controls Avoidance Behavior in Caenorhabditis elegans. Cell Rep. 20, 2501–2512 (2017).

20. Ooi, F. K. & Prahlad, V. Olfactory experience primes the heat shock transcription factor HSF-1 to enhance the expression of molecular chaperones in C. elegans. Sci. Signal. 10, eaan4893 (2017).

21. Jin, X., Pokala, N. & Bargmann, C. I. Distinct Circuits for the Formation and Retrieval of an Imprinted Olfactory Memory. Cell 164, 632–643 (2016).

22. Kauffman, A. et al. C. elegans Positive Butanone Learning, Short-term, and Long-term Associative Memory Assays. J. Vis. Exp. (2011). doi:10.3791/2490

23. Greer, E. R., Pérez, C. L., Van Gilst, M. R., Lee, B. H. & Ashrafi, K. Neural and Molecular Dissection of a C. elegans Sensory Circuit that Regulates Fat and Feeding. Cell Metab. 8, 118–131 (2008).

24. Ha, H. et al. Functional Organization of a Neural Network for Aversive Olfactory Learning in Caenorhabditis elegans. Neuron 68, 1173–1186 (2010).

25. Greer, E. L. et al. Members of the H3K4 trimethylation complex regulate lifespan in a germline-dependent manner in C. elegans. Nature 466, 383–387 (2010).

26. Shirayama, M. et al. piRNAs Initiate an Epigenetic Memory of Nonself RNA in the C. elegans Germline. Cell 150, 65–77 (2012).

27. Troemel, E. R. et al. p38 MAPK Regulates Expression of Immune Response Genes and Contributes to Longevity in C. elegans. PLoS Genet. 2, e183 (2006).

